# Effectors and potential targets selectively upregulated in human KRAS-mutant lung adenocarcinomas

**DOI:** 10.1101/041137

**Authors:** Jinyu Li, Raffaella Sordella, Scott Powers

## Abstract

Genetic and proteomic analysis of human tumor samples can provide an important compliment to information obtained from model systems. Here we examined protein and gene expression from the Cancer Genome and Proteome Atlases (TCGA and TCPA) to characterize proteins and protein-coding genes that are selectively upregulated in *KRAS*-mutant lung adenocarcinomas. Phosphoprotein activation of several MAPK signaling components was considerably stronger in *KRAS*-mutants than any other group of tumors, even those with activating mutations in receptor tyrosine kinases (RTKs) and BRAF. Co-occurring mutations in *KRAS*-mutants were associated with differential activation of PDK1 and PKC-alpha. Genes showing strong activation in RNA-seq data included negative regulators of RTK/RAF/MAPK signaling along with potential oncogenic effectors including activators of Rac and Rho proteins and the receptor protein-tyrosine phosphatase genes PTPRM and PTPRE. These results corroborate RAF/MAPK signaling as an important therapeutic target in *KRAS*-mutant lung adenocarcinomas and pinpoint new potential targets.

## Introduction

How mutationally activated *KRAS* and other canonical *RAS* genes malignantly transform cells and how to block this process for therapeutic benefit has been a subject of intense investigation for over thirty years. The majority of efforts to date have relied on model systems, using established cell lines and mouse models. These studies have identified signaling pathways that are directly stimulated by biochemically active Ras proteins, including the Raf/MAPK and PI3K/Akt pathways^1^. They also have identified pathways or processes further downstream from Ras proteins that are involved in malignant phenotypes induced by mutant *RAS* genes, including the NF–κB pathway^2, 3^, transcriptional activity of the oncogene *YAP*^4^, generation of reactive oxygen species^5^, and anabolic glucose metabolism^6^. A third class of proposed oncogenic mediators of mutant *RAS* genes are induced secreted proteins including TGF–α^7^, Vegf^8^, IL-8^9^, IL-6^10^, CXCL1^11^, and CCL5^12^. Several members of these three classes of mediators of oncogenic Ras have been explored as potential therapeutic targets but as of yet there hasn’t been a clinically successful treatment developed for cancers with mutant *RAS* genes.

The recent completion of large-scale human cancer sample characterizations such as the Cancer Genome Atlas (TCGA) and the Cancer Proteome Atlas (TCPA) has enabled an altogether different approach for discovery of protein targets that are upregulated by mutant *RAS* genes. This approach uses a direct comparison of tumor samples containing mutant *RAS* genes with either corresponding normal tissue samples or with tumor samples that contain wild-type *RAS* genes. One chief advantage of this approach is the analysis is done using the correct *in vivo* physiological context, whereas with model systems there is no guarantee that the physiological context is correct although it is thought that mouse models are superior to cell culture models^13, 14^. Another advantage is that direct analysis of human cancer takes into account the enormous diversity of co-occurring genomic alterations. This diversity requires very large panels of human cancer cell lines to be representative and presents very difficult challenges for mouse modeling.

Recently, three discrete subtypes of human *KRAS*-mutant lung adenocarcinomas were discovered in a breakthrough of our understanding of the variability within this type of lung cancer. These three subtypes have both distinct RNA expression profiles and different patterns of co-occurring mutations in *TP53, STK11, KEAP1*, and others^15^. Furthermore, based on experiments with appropriate cell lines, there appears to be consistent differences in responses to HSP90-inhibitors, suggesting that this classification scheme could be useful in guiding treatment strategies^15^.

In this study, we analyzed genomic and proteomic data from human lung adenocarcinomas to examine the strength and variability of mutant-*KRAS* activation of proteins and genes. *KRAS*-mutants were compared to other lung adenocarcinoma samples and were also examined for the effects of commonly co-occurring mutations.

## Results

### Raf/MAPK signaling proteins are selectively activated in KRAS-mutant lung adenocarcinomas

230 lung adenocarcinoma samples have been comprehensively characterized for mutations and gene expression by TCGA and these same samples have been characterized by reverse-phase protein array (RPPA) for their levels of 160 different proteins and modified proteins^16^. To determine the selective effects of mutational activation of *KRAS* on these proteins in lung adenocarcinomas, we compared *KRAS*-mutant tumors (n= 75) to other tumors with wild-type *KRAS*. We further divided tumors with wild-type *KRAS* into those with other mutations in components of the mitogenic RTK/RAF/MAPK pathway and those without, using the criteria found in the TCGA publication^16^. The other activating mutations in the RTK/RAF/MAPK pathway strongly tended toward mutual exclusivity and included missense or inframe deletion mutations in *EGFR* (n = 26), *ERBB2* (n = 4), or *RIT1* (n = 5); inactivating mutations in *NF1* (n = 19); missense mutations in *BRAF* (n = 16), *MAP2K1* (n = 2), *HRAS* (n = 1), or *NRAS* (n = 1); exon 14 skipping mutations in *MET* (n = 10), gene fusions involving *ROS1* (n = 4), *ALK* (n = 3), or *RET* (n = 2); and focal high-level amplification of *ERBB2* (n = 5), or *MET* (n = 5).

We performed pairwise analysis of the three tumor subsets: KRAS mutants, other RTK/RAF/MAPK mutants, and all others. Since wanted to examine the differences with the greatest potential biological impact, we ranked the differences based on the effect size, rather than *p*–value which can over emphasize very small changes with little variation. We used Cohen’s *d* statistic that is highly related to the signal-to-noise statistic used in gene expression studies^17, 18^. Of all the 160 measured modified proteins and native proteins, the top ranking change in *KRAS*-mutant tumors when compared to either group of wild-type *KRAS* tumors is activation of MEK1, as judged by increased levels detected by the anti-phospho-serine 218,222 MEK1 antibody used by TCPA (Figure 1A). Also top-ranking in both comparisons is phosphorylation activation of MAPK (Figure 1B) and a direct kinase target of MAPK, p90-S6-RSK (Figure 1C). Significant large effects on activation of other direct kinase targets of MAPK (YB-1) and further downstream targets (S6) were also observed (Supplementary Table 1). Also increased in both comparisons of *KRAS*–mutant tumors was activation of mTOR as determined by phosphorylation at serine 2448 (Figure 1D). However, both PDK1 and AKT kinases, which might be expected to be upregulated in *KRAS*–mutant lung cancers due to Kras protein interaction with PI3-kinase, did not show any significant phosphorylation activation in *KRAS*–mutant tumors (Supplementary Table 1).

Analysis of a well-validated mouse model of *KRAS*–mutant lung adenocarcinoma revealed that tumors showed significant activation of NF-κb and furthermore that these tumors were dependent upon NF-κb activity for tumor maintenance^3^. We did not observe any activation of NF-κb in human *KRAS*-mutant lung adenocarcinomas; in fact, NF-κb activation was significantly lower in KRAS-mutant tumors compared to other Raf/MAPK mutant tumors (Supplementary Table 1).

### Other mutations affecting the RTK/RAF/MAPK pathway have specific effects on protein levels

We next extended our comparative protein analysis to look within the group with other mutations in the RTK/RAF/MAPK pathway. The subgroups with sufficiently large numbers for statistical analysis included *EGFR* mutants (n = 26), *NF1* mutants (n = 19), *MET* mutants (n = 15), and *BRAF* mutants (n = 16). There were significant differences (Supplementary Table 2), the most prominent of which are highlighted in Figure 2. Notably, the phosphorylation activation of EGFR and HER2 proteins was significantly higher in *EGFR* mutants, but there was also increased phosphorylation of RET protein and decreased phosphorylation of HER3 and MET proteins (Figure 2). *MET* mutants on the other hand showed opposite effects, with significant increased phosphorylation of MET protein but decreased phosphorylation of EGFR, HER2, and RET (Figure 2). *NF1* mutants also showed decreased phosphorylation of EGFR, HER2, and RET proteins (Figure 2; Supplementary Table 2). We did not observe significant effects with these select phosphoproteins in *BRAF* mutants (Supplementary Table 2).

### Effect of co-occurring mutations on protein levels in KRAS-mutant lung adenocarcinomas

We next examined the effects of co-occurring mutations in *TP53, STK11*, and *KEAP1* on protein levels in mutant-*KRAS* tumors. Using cBioPortal analysis of TCGA lung adenocarcinomas, we confirmed that mutations in *TP53* and *STK11* tend to be mutually exclusive, whereas mutations in *KEAP1* and *STK11* tend to co-occur. Amongst *KRAS*-mutant tumors, there were 22 tumors with mutations in *TP53* but not *STK11* or *KEAP1*, 20 tumors with mutations in *STK11* but not in *TP53*, and 13 tumors with mutations in *KEAP1* but not in *TP53* (8 of these tumors also had mutations in *STK11*). There was only one *KRAS*-mutant tumor with mutations in both *TP53* and *STK11* or *KEAP1*. Excluding this single sample, we divided *KRAS*–mutant tumors into three groups: those with *TP53* mutations (n = 20), those with *STK11* and/or *KEAP1* mutations (n = 25), and those without any of these mutations (n = 27). Pairwise analysis of these three groups revealed that the phosphorylation activation status of Raf/MAPK pathway components was not significantly affected (Supplementary Table 3).

The strongest effect of co-occurring *TP53* mutations was seen with increased levels of Annexin I protein (Figure 3A). As expected, the group with mutations in the protein kinase gene *STK11* had significantly lower levels of activation of its direct target, AMPK, than the other two groups (Figure 3B). More surprising however were the significantly lower levels of phosphorylation activated PDK1 and PKC-alpha proteins in this same group (Figure 3C and 3D).

### Top ranked KRAS-mutant induced genes by RNA-seq data

We then looked at RNA-seq data to find genes with the greatest induction specifically in *KRAS*-mutant lung adenocarcinomas. We ranked genes based on the average effect size when *KRAS*-mutant tumors were compared to tumors with other mutations in genes of the Raf/MAPK pathway, and when compared to tumors with no mutations in genes of the Raf/MAPK pathway. *KRAS* itself was the top-ranked induced gene (Figure 4; Supplementary Table 4). Within the top-100 induced genes were five negative regulators of RTK/MAPK signaling (*DUSP4, DUSP6, NF1, SPRED2, SPRY4*); the canonical Raf/MAPK transcriptional targets *ETV4 and ETV5* (ETS-family members) and *FOS* (Figure 4; Supplementary Table 4). Also within this top-ranked group are several candidate transcriptional oncogenic effectors for mutant *KRAS*, including the receptor tyrosine kinase genes *INSR* and *IGFR1*; the receptor tyrosine phosphatase genes *PTPRE* and *PTPRM*; and genes encoding guanine-nucleotide exchange-factors that activate Rac/Rho/Cdc42 proteins *DOCK5, DOCK1, DNMBP*, and *PLEKHG2;* (Figure 4). Also included in the top-100 ranked induced genes is the gene *CXorf61* which encodes a tumor antigen termed Kita-Kyushu lung cancer antigen 1 (Figure 4).

Approximately 50% of the aforementioned genes that are selectively induced in *KRAS*-mutant lung adenocarcinomas were also significantly affected by co-occurring mutations in *TP53, STK11*, or *KEAP1*. For the most part, co-occurring mutations in *STK11* and/or *KEAP1* were associated with significantly stronger expression of these genes. In only one case(*DUSP6*) was co-occurrence of mutant *TP53* associated with stronger expression (Figure 5).

## Discussion

Several components of the Raf/MAPK pathway show strong selective activation in *KRAS*-mutant lung adenocarcinomas, providing further corroboration that this pathway is a key therapeutic target for *KRAS*-mutant lung tumors. However, a potentially important finding of this study is that many of the signaling proteins and pathways thought to be activated in *KRAS*–mutant human lung adenocarcinomas based on studies with models systems are not activated, at least not at steady state levels as assayed by immunoblotting of tissue samples. This makes it less likely that these pathways or proteins correspond to *KRAS*-selective dependencies. These proteins include PI3-kinase (as judged by activation of AKT), AKT, and NF-κB, all of which have been proposed to be important mediators of mutant *KRAS* in lung adenocarcinomas^3, 19^. Since mTOR is the only component of the PI3-kinase pathway that was selectively activated in *KRAS*-mutant tumors, a greater therapeutic window could conceivably be achieved by combining Raf/MAPK inhibitors with highly selective mTOR inhibitors that do not inhibit PI3-kinase.

All three subtypes of *KRAS*-mutants showed approximately equivalent activation of Raf/MAPK components. However, there were significant differences in protein and phosphoprotein levels detected in the three subgroups. The subgroup comprised of *KRAS/STK11* and *KRAS/KEAP1* double mutants, along with *KRAS/STK11/KEAP1* triple mutants, had significantly lowers levels of phosphorylated AMPK, which is to be expected since AMPK is a direct kinase target of STK11. However this group was more unexpectedly associated with lower levels of phosphorylation activation of PDK1 and PKC-alpha, both of which have been actively pursued targets for developing potentially clinically effective inhibitors. However in this case, it would appear that loss of STK11 function is likely driving this decrease in activity, and that as such this would not represent a potential selective dependency in any of the *KRAS*-mutant subgroups.

Genomic studies usually rank differentially expressed genes by p-value or FDR. However, this ranking method does not taking into account the size of the effect. Potential transcriptional effectors of mutant-*KRAS* would more likely be genes showing the largest consistent increase in RNA. Therefore we used Cohen’s *d* statistic, closely related to the signal-to-noise statistic, to rank genes selectively upregulated in *KRAS*-mutant tumors. Amongst the top 100 upregulated genes were several negative regulators of Raf/MAPK signaling, consistent with an important known aspect of negative feedback regulation in this pathway^20, 21^, as well as canonical transcriptional targets for Raf/MAPK signaling including *FOS* and the ETS-family members *ETV4* and *ETV5*. Additionally, several potential oncogenic transcriptionally activated targets were uncovered including *IGF1R* and *INSR*. A connection between mutant *KRAS* and IGF1R in lung adenocarcionoma has previously been documented^22^. We detected transcriptional activation of several guanine-nucleotide exchange factors that activate different proteins in the Rac/Rho/Cdc42 family. Thus the effect of mutational activation of KRAS on guanine-nucleotide exchange factors for small GTP-binding proteins is broader in lung adenocarcinoma than its biochemical activation of exchange factors that activate Ral proteins. Upregulation of the receptor protein-tyrosine phosphatase gene *PTPRE*, the third highest ranked upregulated gene, is of particular interest since it has been shown to be upregulated by RAS in a mouse model of mammary carcinoma and to possess on its own the ability to promote mammary tumor formation^23, 24^. Finally, this set of upregulated genes includes a potential immunotherapy target for *KRAS*-mutant lung adenocarcinomas, the Kita-Kyushu lung cancer antigen 1, encoded by the *CXorf61* gene^25^. This protein has also been shown to be a potential target for T cell based therapies in triple-negative breast cancer^26^.

**Figure legends**

**Figure 1.**
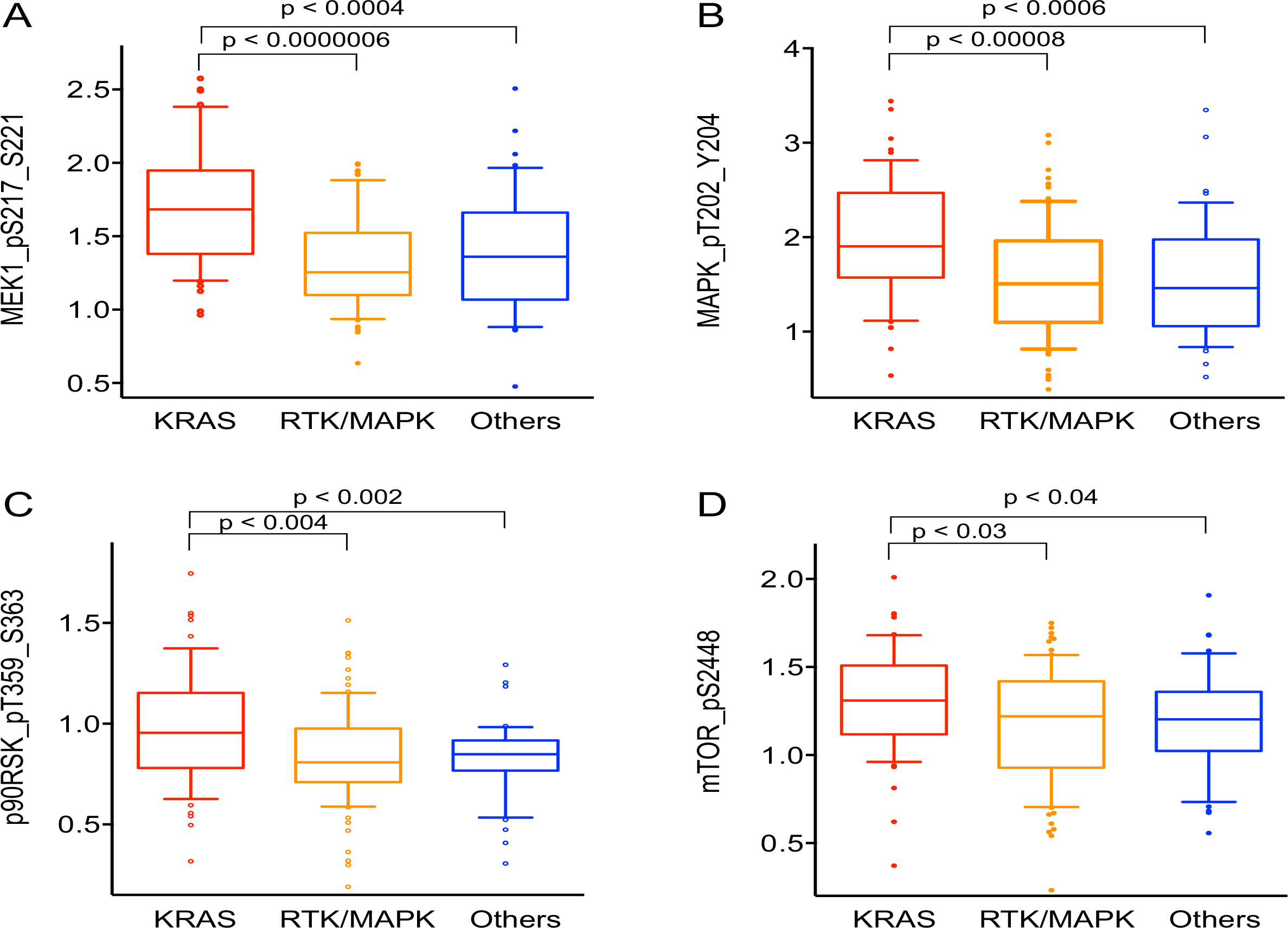
Relative levels of select phosphoproteins in three subgroups of human lung adenocarcinomas. All values are plotted on an arbitrary log2 scale to highlight relative levels rather than absolute amounts. For each panel, the red colored group represents *KRAS*-mutant tumors, the orange colored group represents other Raf/MAPK pathway mutants, and the blue colored group represents all other tumors. Brackets with associated p-values indicate significant differences. (A) levels of MEK1_pS217_S221; (B) levels of MAPK_pT202_Y204; (C) levels of p90RSK_pT359_S363; (D) levels of mTOR_pS2448.

**Figure 2.**
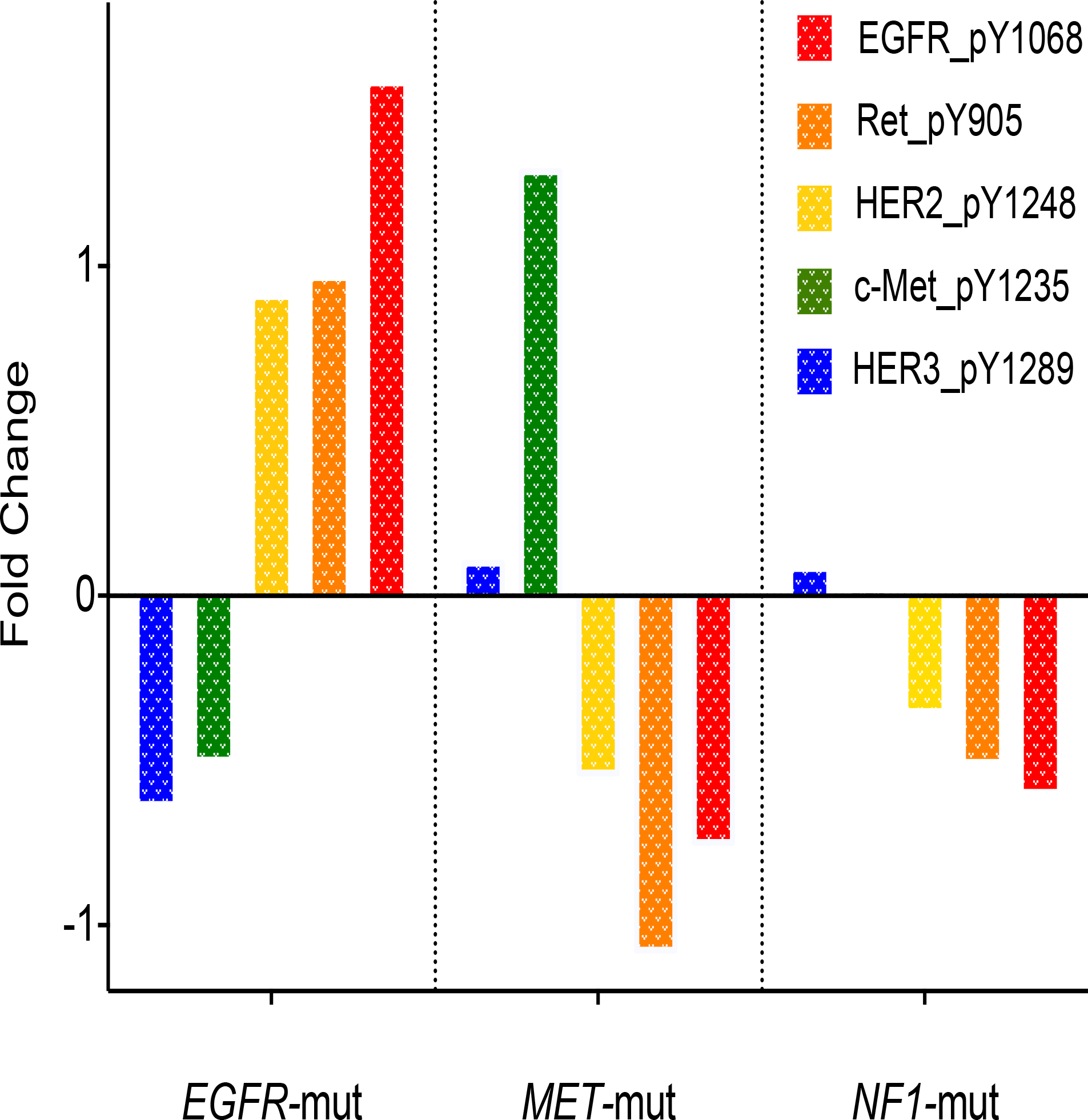
Relative levels of select phosphoproteins within *KRAS*-wild type lung adenocarcinomas with other mutations in the RTK/RAF/MAPK pathway. Pairwise comparisons were made between *EGFR* mutants (n = 29) and all other mutants (n = 71); *NF1* mutants (n = 19) and all other mutants (n = 81); *MET* mutants (n = 15) and all other mutants (n = 85).

**Figure 3.**
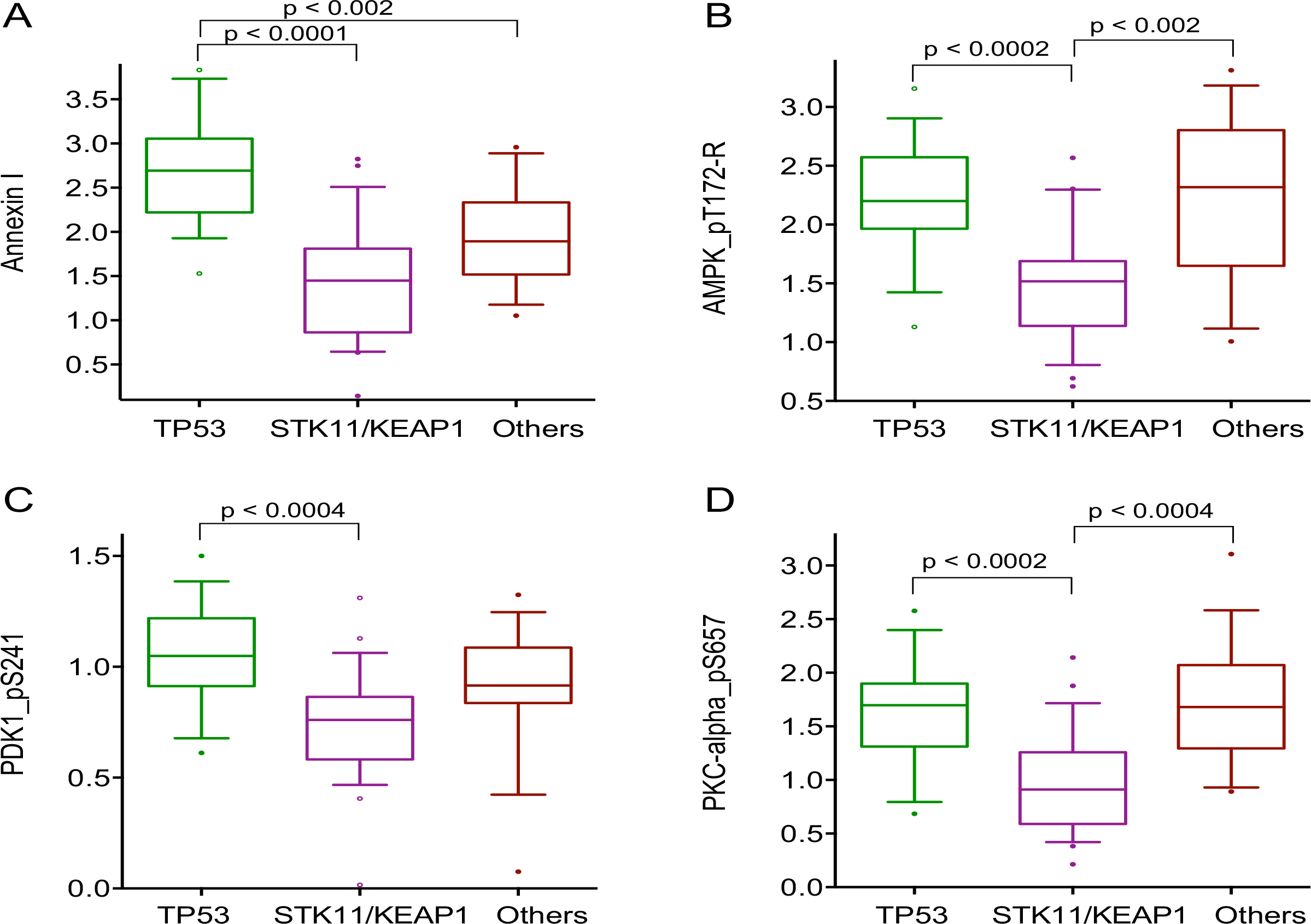
Relative levels of select proteins phosphoproteins in three subgroups of KRAS-mutant adenocarcinomas. All values are plotted on an arbitrary log2 scale to highlight relative levels rather than absolute amounts. For each panel, the green colored group represents *KRAS/TP53* doublemutant tumors, the violet colored group represents other *KRAS/STK11, KRAS/KEAP1*, and *KRAS/STK11/KEAP1* compound mutants, and the brown colored groups *KRAS* mutants without those other mutations. Brackets with associated p-values indicate significant differences. (A) levels of Annexin I; (B) levels of AMPK_pT172-R; (C) levels of PDK1_pS241; (D) levels of PKC-alpha_pS657.

**Figure 4.**
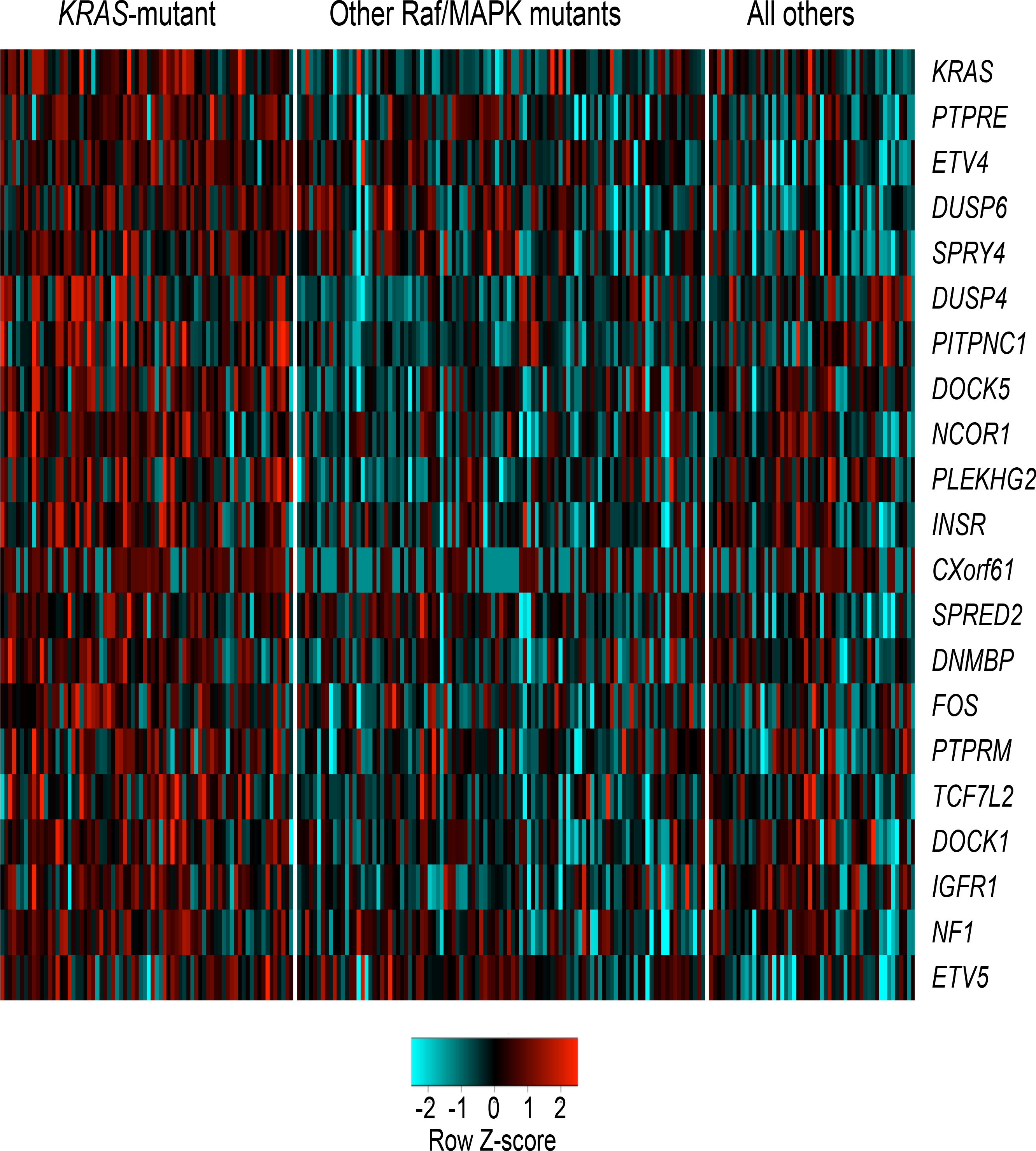
Heatmap showing the relative RNA values of select genes in three subgroups of human lung adenocarcinomas. The three subgroups include *KRAS*-mutant tumors, other Raf/MAPK pathway mutants, and all other tumors. The select genes are indicated on the right. Row values were normalized and scaled and presented as Z-scores.

**Figure 5.**
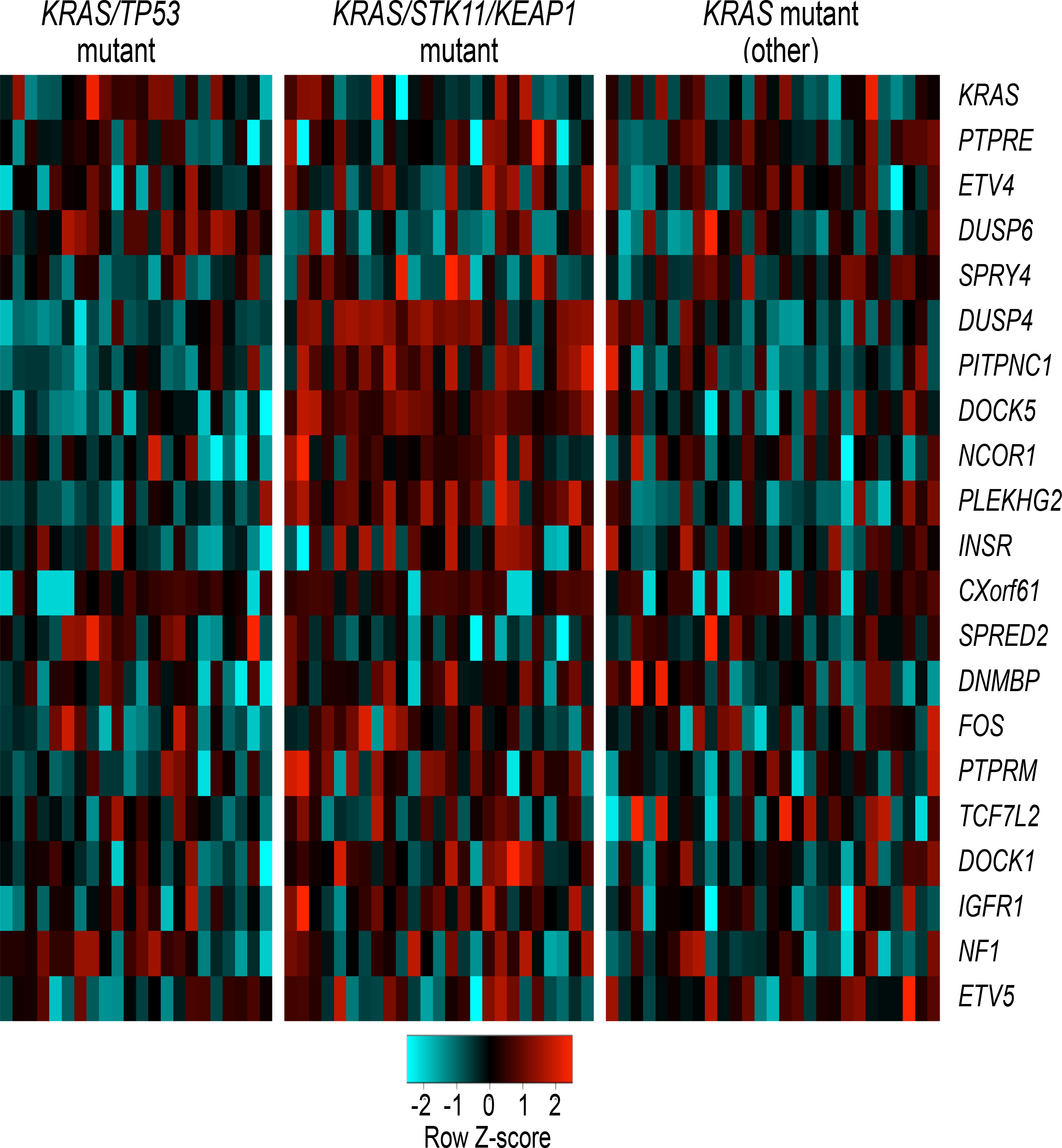
Heatmap showing the relative RNA values of select genes in three subgroups of *KRAS*-mutant human lung adenocarcinomas. The three subgroups are *KRAS/TP53* double mutants; *KRAS/STK11, KRAS/KEAP1*, and *KRAS/STK11/KEAP1* compound mutants; and *KRAS* mutants without those other mutations. The select genes are indicated on the right. Row values were normalized and scaled and presented as Z-scores.

## Methods

We used mutational, RNAseq and protein data in our analysis from the TCGA lung adenocarcinoma project. All data downloaded was Level 3. Mutation data was downloaded from cBioPortal for Cancer Genomics (www.cbioportal.org), under Lung Adenocarcinoma (TCGA, Provisional) with “All Complete Tumors (230)”. RNAseq data was downloaded from FireBrowse by Broad Institute (www.firebrowse.org), choosing “illuminahiseq_rnaseqv2-RSEM_genes”. Protein data was downloaded from The Cancer Proteome Atlas (TCPA) project in MD Anderson Cancer Center (bioinformatics.mdanderson.org/main/TCPA). To generate heatmaps, the R package “gplots” was used.

We segregated samples using cBioPortal into three mutually exclusive groups: ones with *KRAS* missense mutations, ones with mutations in genes encoding other components of the RTK/RAF/MAPK pathway as delineated in the text, and other samples. Those three subgroups were used to perform pairwise comparisons using protein (RPPA) and RNAseq data. We used Cohen’s d to measure effect size using the R-package “effsize”. Ranking with both effect size and t-test p-value generated top-ranked proteins and genes induced in *KRAS* mutants or Raf/MAPK pathway mutants (Figure 1, Supplementary Table 1). We used a similar approach to divide *KRAS* mutants into three groups depending on *TP53, STK11*, or *KEAP1* status.

## Acknowledgement

This work was supported in part by NIH grant U01CA168409.

## Author Contributions

Conceived and designed the analysis: SP. Computational analysis: JL. Analyzed data: JL, SP, RS. Wrote the paper: JL, SP.

## Additional Information

The authors have declared that no competing financial interests exist.

## Supplementary Information

**Supplementary Table Legends**

**Supplementary Table 1.** Protein levels in three groups of lung adenocarcinomas. Mean levels in three groups are listed along with the p-values and effect sizes of pairwise comparisons. G1 = *KRAS*-mutants; G2 = other Raf/MAPK pathway mutants; G3 = all others.

**Supplementary Table 2.** Relative protein levels in four groups of *KRAS*-wild-type lung adenocarcinomas that have other mutations in the RTK/RAF/MAPK pathway. p-values in the four groups are listed along with the effect sizes of pairwise comparisons.

**Supplementary Table 3.** Protein levels in three groups of *KRAS*-mutant lung adenocarcinomas. Mean levels in three groups are listed along with the p-values and effect sizes of pairwise comparisons. G1 = *KRAS/TP53* double mutants; G2 = *KRAS/STK11* or *KRAS/KEAP1* double mutants and *KRAS/STK11/KEAP1* triple mutants; G3 = all others.

**Supplementary Table 4.** RNA levels in three groups of lung adenocarcinomas. Mean levels in three groups are listed along with the p-values and effect sizes of pairwise comparisons. G1 = *KRAS*-mutants; G2 = other Raf/MAPK pathway mutants; G3 = all others.

